# Recurrent chromosome reshuffling and the evolution of neo-sex chromosomes in parrots

**DOI:** 10.1101/2021.03.08.434498

**Authors:** Zhen Huang, Ivanete Furo, Valentina Peona, Jing Liu, Anderson J. B. Gomes, Wan Cen, Hao Huang, Yanding Zhang, Duo Chen, Xue Ting, Youling Chen, Qiujin Zhang, Zhicao Yue, Alexander Suh, Edivaldo H. C. de Oliveira, Luohao Xu

## Abstract

The karyotype of most birds has remained considerably stable during more than 100 million years’ evolution, except for some groups, such as parrots. The evolutionary processes and underlying genetic mechanism of chromosomal rearrangements in parrots, however, are poorly understood. Here, using chromosome-level assemblies of three parrot genomes (monk parakeet, blue-fronted amazon, budgerigar), we uncovered frequent chromosome fusions and fissions among parrots, with most of them being lineage-specific. In particular, at least 12 chromosomes recurrently experienced inter-chromosomal fusions in different parrot lineages. Two conserved vertebrate genes, *ALC1* and *PARP3,* with known functions in the repair of double-strand breaks and maintenance of genome stability, were specifically lost in parrots. The loss of *ALC1* was associated with multiple deletions and an accumulation of CR1-psi, a novel subfamily of transposable elements (TEs) that recently amplified in parrots, while the loss of *PARP3* was associated with an inversion. Additionally, the fusion of the ZW sex chromosomes and chromosome 11 has created a pair of neo-sex chromosomes in the ancestor of parrots, and the chromosome 25 has been further added to the sex chromosomes in monk parakeet. The newly formed neo-sex chromosomes were validated by our chromosomal painting, genomic and phylogenetic analyses. Transcriptome profiling for multiple tissues of males and females did not reveal signals of female-specific selection driving the formation of neo-sex chromosomes. Finally, we identified one W-specific satellite repeat that contributed to the unusual enlargement of the W chromosome in monk parakeet. Together, the combination of our genomic and cytogenetic analyses highlight the role of TEs and genetic drift in promoting chromosome rearrangements, gene loss and the evolution of neo-sex chromosome in parrots.

## Introduction

The karyotypes and genome sizes of birds have remained considerably stable over more than 100 million years’ evolution of modern birds (Zhang et al. 2014; Ellegren 2010; Kretschmer, Ferguson-Smith, and de Oliveira 2018; Kapusta, Suh, and Feschotte 2017; Damas et al. 2019). A typical avian karyotype consists of about 40 pairs of chromosomes (2n = 80), among which about 30 paris are microchromosomes (smaller than 20 Mb). Among the 12% of bird species with documented karyotype, most have diploid numbers ranging from 76 to 82 (Kretschmer, Ferguson-Smith, and de Oliveira 2018; Zhang 2018; Kapusta and Suh 2017). Both cytogenetic mapping and genome assembly have revealed the deep conservation in synteny of both macrochromosomes (larger than 20 Mb) and microchromosomes (O’Connor et al. 2019; Damas et al. 2018; O’Connor, Romanov, et al. 2018; Romanov et al. 2014; Peona et al. 2021; Liu et al. 2021; Li et al. 2021). For instance, emu (a deep-branching bird species) and chicken differ only by one single chromosomal fusion event since their divergence about 100 million years ago (Liu et al. 2021; Shetty, Griffin, and Graves 1999).

Despite the overall conservation, the variation of karyotype is apparent in some bird lineages (O’Connor, Farré, et al. 2018; Kretschmer, Ferguson-Smith, and de Oliveira 2018). Many raptor species (birds of prey) have much fewer chromosomes, and particularly, some falcons have a haploid number as low as 20 (Joseph et al. 2018; Oliveira et al. 2005; de Boer 1975). Psittaciformes (parrots, macaws, and alleys) is another bird lineage displaying pronounced karyotype variation (Francisco and Galetti Júnior 2001; Rus et al. 2016; Furo et al. 2020). For instance, our previous cytogenetic work characterizing the karyotype of monk parakeet (*Myiopsitta monachus*) and blue-fronted amazon (*Amazona aestiva*) revealed their diploid numbers of 48 and 70, respectively (Furo et al. 2017), suggesting that chromosome rearrangements in birds can occur rapidly within a short period of time. It is unclear, however, how and why some bird groups have more frequent chromosome rearrangements than others.

It is also unclear how often those chromosomal rearrangements may involve sex chromosomes. The fusion of the sex chromosome and a pair of autosomes can create what is called a ‘neo-sex chromosome’. Neo-sex chromosomes have been frequently reported in various organisms, including muntjac deer (Zhou et al. 2008), threespine sticklebacks (Ross et al. 2009; Kitano et al. 2009), *Drosophila obscura* group species (R. Bracewell and Bachtrog 2020), monarch butterflies (Mongue et al. 2017), parasitic nematodes (Foster et al. 2020), among other animals (Jetybayev et al. 2017). Studies across many taxa with neo-XY chromosomes reveal adaptive evolution of the neo-Y chromosomes that plays a role in resolving sexual antagonism and speciation (Kitano et al. 2009; Zhou and Bachtrog 2012; R. R. Bracewell et al. 2017). In birds, the sex chromosome system is female-heterogametic (ZW female, ZZ males) and is highly conserved across bird lineages (Xu and Zhou 2020; Zhou et al. 2014; Xu, Wa Sin, et al. 2019). While the presence of neo-sex chromosomes, to date, has been shown in a few songbirds (Sigeman, Ponnikas, and Hansson 2020; Gan et al. 2019; Sigeman et al. 2019; Pala et al. 2012; Dierickx et al. 2020) and recently a cuckoo (Kretschmer et al. 2020), it remains unclear whether neo-sex chromosomes have evolved in parrots.

In this study, we produced chromosome-level assemblies of monk parakeet and blue-fronted amazon genomes, and re-produced the chromosome-level assembly of budgerigar (*Melopsittacus undulatus*). Our comparative analysis reveals the dynamic evolutionary history of parrot karyotypes which were shaped by frequent and independent inter-chromosomal fusions and fissions. We also discovered and characterized neo-sex chromosomes in parrots, and investigated the evolutionary consequence of the sex chromosome fusions. Finally, we identified one satellite sequence that was likely responsible for the enlargement of the W chromosome in the monk parakeet.

## Results

### Chromosome-level genome assembly of three parrots

We used chromatin conformation capture (Hi-C) sequencing data to produce chromosome-level assemblies based on a new long-read genome of the monk parakeet generated in this study and two existing draft genomes of the blue-fronted amazon (Wirthlin et al. 2018) and the budgerigar (Ganapathy et al. 2014) (**Table 1**). More than 97.5% of contig sequences of monk parakeet were scaffolded into 24 chromosome models (**Supplementary Fig. S1**), consistent with the known haploid chromosome number (Furo et al. 2017). Among the unanchored contigs (25.8 Mb in length), 89.5% of the sequences are tandem repeats or TEs, suggesting very few non-repetitive genomic sequences are missing from the chromosome assembly. This female genome of monk parakeet also contains a W chromosome that is 13.8 Mb long, rendering it among the largest assembled avian W chromosomes to date (Peona et al. 2021, 2020). Moreover, we identified a candidate centromeric satellite sequence which is 191 bp long and is validated by the fluorescent *in situ* hybridization (FISH) experiment (**Supplementary Fig. S2**). For the short-read-based draft genomes of blue-fronted amazon (Wirthlin et al. 2018) and budgerigar (Ganapathy et al. 2014), we anchored 99.5% and 97.6% of the assembled sequences into 29 and 22 chromosome models, respectively. According to the known karyotype of blue-fronted amazon (2n = 70) (Furo et al. 2017) and budgerigar (2n = 62) (Nanda et al. 2007), chromosome models of six and nine presumably small microchromosomes, were likely not scaffolded in the respective chromosome assemblies.

**Table 1.** Genome assemblies of three parrots.

|  | Monk parakeet | blue-fronted amazon | budgerigar |
| --- | --- | --- | --- |
| Genome size (Gb) | 1.17 | 1.13 | 1.09 |
| N50 contig (Mb) | 24.5 | 0.4 | 0.05 |
| N50 scaffold (Mb) | 75.7 | 89.0 | 101.4 |
| % assigned to chromosomes | 97.8 | 99.5 | 97.6 |
| BUSCO complete (%) | 94.1 | 93.3 | 93.6 |
| Repeat content (%) | 14.6 | 9.5 | 9.7 |

To estimate the divergence time among parrot species, we included the genomes of two other parrots (sun parakeet (Gelabert et al. 2020) and kea (Zhang et al. 2014)), two songbirds (Laine et al. 2016; Peona et al. 2021) and two outgroups (chicken and emu) (Liu et al. 2021; Warren et al. 2017). The phylogenetic tree built with 1.4 million 4-fold degenerate sites is consistent with previous knowledge of parrot phylogeny (Gelabert et al. 2020; Wright et al. 2008; Oliveros et al. 2019), and dated the common ancestor of parrots to about 37.2 million years ago (**Fig. 1a**). The pair of monk parakeet and blue-fronted amazon whose haploid numbers differ by 11 shared a common ancestor 19.9 million years ago (**Fig. 1a**).

**Fig. 1.**
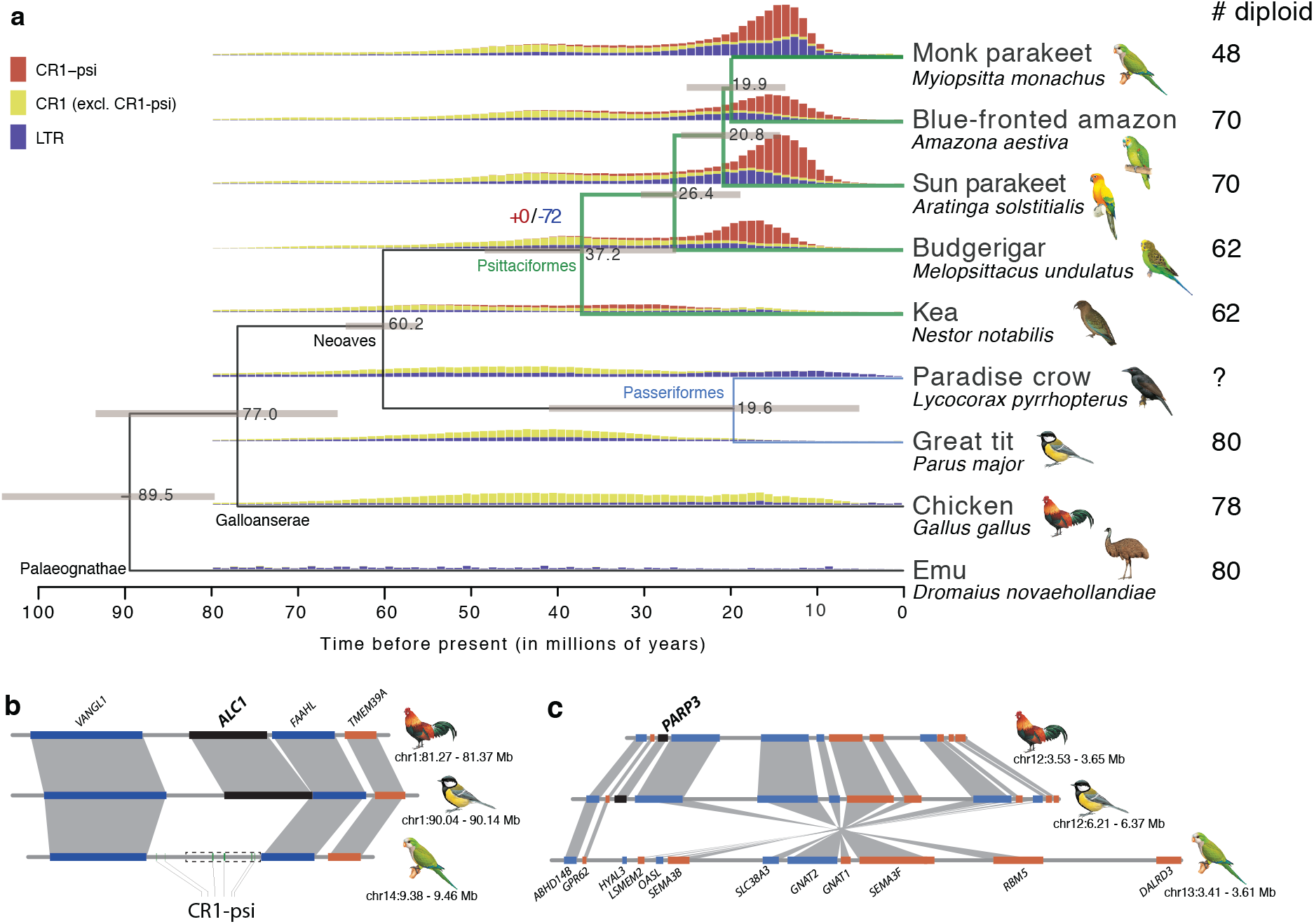
Phylogeny and comparative genomics of parrots. **a)** The phylogeny of nine bird species shows the divergence time (denoted at the nodes) calibrated with fossil records at the node Psittacopasserae (56-64.5 MY) and Aves (80-105 MY) (Oliveros et al. 2019). The numbers of gene gains (red) and losses (blue) were denoted at the node of Psittaciformes (parrots). The vertical bars show the timing of CR1 and LTR retrotransposon insertions, with the height representing the frequency of TE activity. The numbers of diploid chromosomes are listed in the right panel. **b-c)** The loss of *ALC1* and *PARP3* (highlighted in black) in parrots. Monk parakeet is used to represent the parrot lineage. The grey bands indicate the connection of orthologs across species. *ALC1* was pseudogenized (indicated by dashed rectangle) due to multiple exon losses (Supplementary Fig. S6). Multiple copies of CR1-psi have been inserted at the *ALC1* locus. *PARP3* is located at the breakpoint of a parrot-specific inversion. Illustrations reproduced by permission of Lynx Edicions.

### Expansion of a novel TE family in parrots

The proportion of transposable elements (including LTR and non-LTR retrotransposons) in parrot genomes (mean 9.6%) is slightly higher than that in songbirds (~7.8%, the sister group of parrots) or most other birds (Feng et al. 2020), mainly due to the increased activity of chicken repeat 1 (CR1) non-LTR retrotransposons (**Supplementary fig. S3a**). We identified a parrot-specific subfamily of the CR1-E family, named CR1-psi, accounting for about half of the parrot CR1 content (**Fig. 1a**, **Supplementary fig. S3b-c**). While most of the CR1-psi copies were severely 5’ truncated, we detected on average 1,860 larger copies of CR1-psi (i.e., >2 kb) that are evolutionarily young and tend to be clustered by parrot species, suggesting their recent lineage-specific propagation (**Fig. 1a**, **Supplementary fig. S4**).

### Parrot-specific gene loss

To examine the changes in gene content of parrot genomes relative to other birds, we further included a non-avian reptile, the green anole (Alföldi et al. 2011), to reconstruct the evolutionary history of bird gene families. While we failed to identify any parrot-specific gene gains, at least 72 genes were found to be lost in all five parrot genomes but present in the other sampled birds (**Fig. 1a**, **Supplementary Table S1**). We further confirmed that those 72 genes were not present in the transcriptomes of nine different tissues of monk parakeet (**Supplementary Table S2**), suggesting they were probably not the hidden genes (Botero-Castro et al. 2017; Yin et al. 2019). We did not detect any functional enrichment for the lost genes, but two of them, *ALC1* and *PARP3*, are related to repair of DNA damage and maintenance of genome stability (Ahel et al. 2009; Day et al. 2017; Mejías-Navarro et al. 2020; Boehler et al. 2011; Beck et al. 2014, 2019; Sellou et al. 2016; Tsuda et al. 2017; Belousova et al. 2018; Zarkovic et al. 2018; Grundy et al. 2016). Among the 72 lost genes, 24 reside in conserved synteny blocks (collinear gene order) across bird species (**Fig. 1b**, **Supplementary fig. S5**), and the gene loss of five of them, including *ALC1*, seems to be associated with CR1-psi. At the locus where *ALC1*, for instance, has been pseudogenized through multiple independent exon deletions (**Supplementary Fig. S6**), we detected a remarkable abundance of partial copies of CR1-psi (**Fig. 1b**). This reflects the nature of mutational hotspot in this region that initially led to pseudogenization of *ALC1*, but it’s also possible that sequence deletion due to non-allelic homologous recombination between CR1-psi copies contributed to exon losses (González and Petrov 2012). We also found 6 lost genes located at the breakpoints of conserved synteny blocks due to rearrangement events in parrots. (**Supplementary Table S3**). For instance, *PARP3* is located at the boundary of a parrot-specific inversion involving 9 genes (**Fig. 1c**). This suggests the potential role of chromosomal rearrangements (inversions) in gene loss (Furuta et al. 2011; Calvete et al. 2012).

### Frequent and independent inter-chromosomal rearrangements

The decreased number of chromosomes in parrots compared to the typical number in avian karyotypes suggests the occurrence of multiple chromosomal fusion events in this lineage. To reconstruct the evolutionary history of chromosomal changes, we aligned the chromosome-level assemblies of three parrots and four outgroups that have chromosome-level assemblies, namely two passerines (Neoaves), chicken (Galloanserae) and emu (Palaeognathae), thus covering all three major bird clades. Between the four non-parrots, typically no more than two events of inter-chromosomal rearrangements can be found in pairwise comparisons, but within the parrot lineage, these comparisons suggest that multiple chromosomal fusions or fissions events occurred independently (**Fig. 2**). We found that only three chromosomal fusions (chr6+chr7, chr8+chr9 and chr11+chrZ) and one fission (chr7) are shared by all parrots (**Fig. 2**, **Supplementary fig. S7**). In addition, none of the chromosomal changes are common to the ancestral lineage leading to monk parakeet and blue-fronted amazon except for a fission of chr2 (**Fig. 2**). In other words, most chromosomal rearrangements are specific to each sampled species and likely have recent origins. Some chromosomal segments have experienced multiple rearrangements in a short time. For instance, an ~18 Mb sequence (chr2A) was split from the ancestral chr2 and further experienced one fusion and one fission in the lineage leading to monk parakeet (**Fig. 2**). Monk parakeet and budgerigar have lower haploid chromosome numbers than blue-fronted amazon (24 and 31 vs. 35), which can be explained by more chromosomal fusions (**Fig. 2**). In particular, we identified eight events in which a microchromosome has fused with a macrochromosome, but have not identified any inter-microchromosome fusion events, in agreement with the suggestion by studies in falcons that reduction of chromosome number is mainly due to fusions of microchromosomes with macrochromosomes (Wilcox, Boissinot, and Idaghdour 2019). In addition, chromosomal fissions likely led to two and three new microchromosomes in monk parakeet and blue-fronted amazon, respectively (**Fig. 2**), but such newly formed microchromosomes seem to be rare in budgerigar or other birds.

**Fig. 2.**
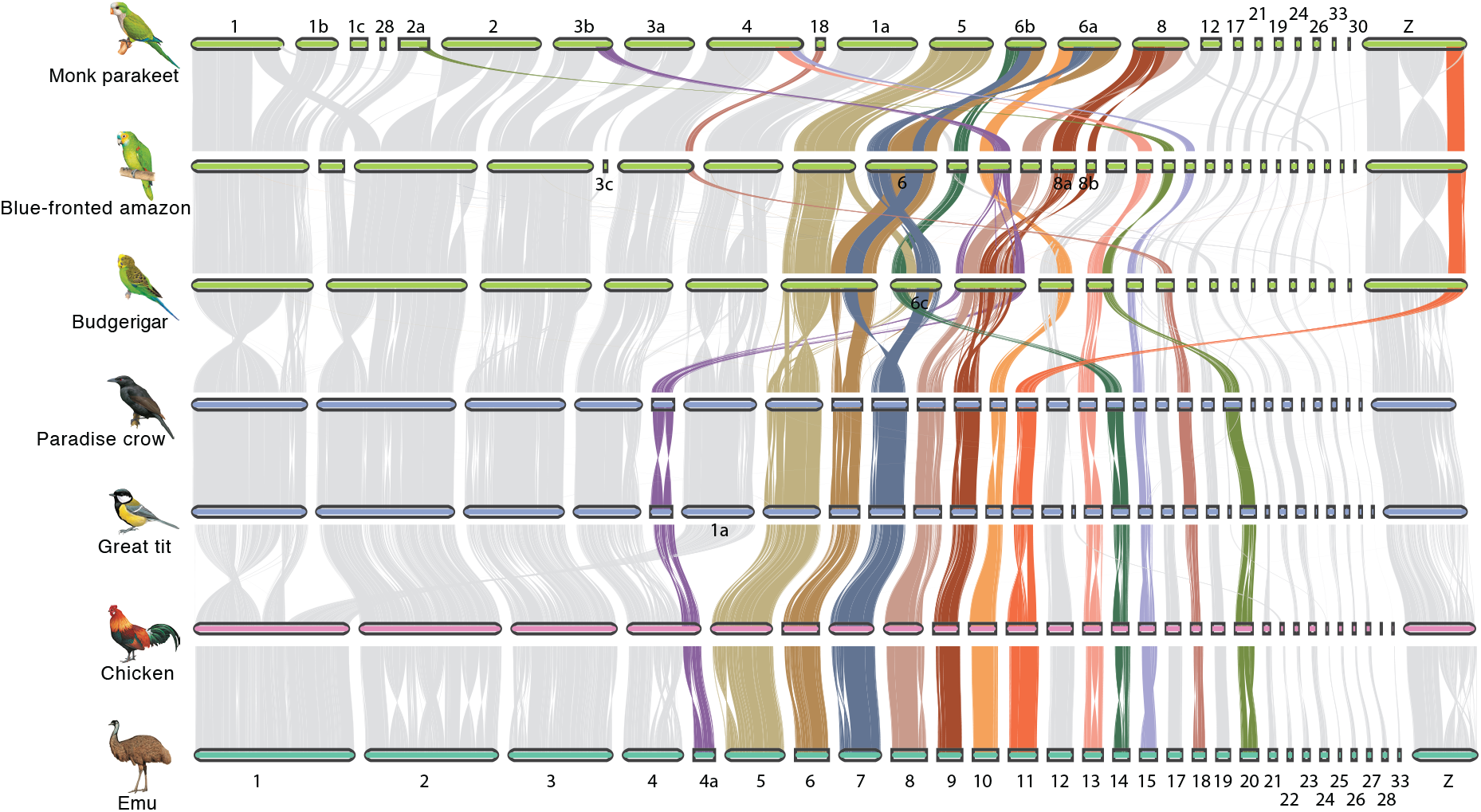
Frequent chromosomal rearrangements across parrot genomes. Pairwise whole-genome alignments across seven chromosome-level assemblies of bird genomes. Each horizontal bar represents one chromosome, and chromosome IDs of chicken were labeled at the bottom; for parrot genomes some chromosomes were renamed following chromosome fissions. The 12 chromosomes that experienced recurrent chromosomal rearrangements are highlighted in colors.

The frequency of chromosomal rearrangements varies among chromosomes, with some chromosomes experiencing repeated and independent rearrangements. We found that 12 chromosomes (chr4A, chr5, chr6, chr7, chr8, chr9, chr10, chr13, chr14, chr15, chr18, chr20) have experienced at least two independent chromosomal changes in parrots. For example, chr10 was fused with chr12 in budgerigar, fused with chr4a in blue-fronted amazon, but fused with chr6a in monk parakeet (**Fig. 2**). Those more frequently rearranged chromosomes tend to have an intermediate chromosome size (20 - 40 Mb) (**Fig. 2**).

### Evolutionary breakpoints of inter-chromosomal rearrangements

We next asked whether the chromosomal changes were adaptive due to, for instance, the creation of a new linkage of formerly unlinked chromatin (Guerrero and Kirkpatrick 2014). Despite independent origins for many of the evolutionary breakpoints (EBPs) of fissions among parrot species, they tend to be located in regions with lower insulation scores in the pre-fission chromosomes (Wilcoxon test, p < 0.05) (**Fig. 3a**), i.e. stronger interacting barriers between the two flanking chromatins. For instance, a fission EBP at 107 Mb on chr1 of blue-fronted amazon has the lowest insulation score along the chromosome, separating two mega-scale flanking chromatin domains (**Fig. 3b**). Since the flanking chromatins separated by this EBP already have infrequent interactions, a breakdown of their physical linkage, as occurred in monk parakeet (**Fig. 3c**), probably has a limited impact in disrupting the *cis*-regulation, if any. Similarly, at the fusion EBPs, the insulation scores tend to be lower, though only significantly so in blue-fronted amazon (Wilcoxon test, p < 0.05) (**Fig. 3d**), suggesting the newly joined chromosomes have not yet established frequent chromatin interactions between them.

**Fig. 3.**
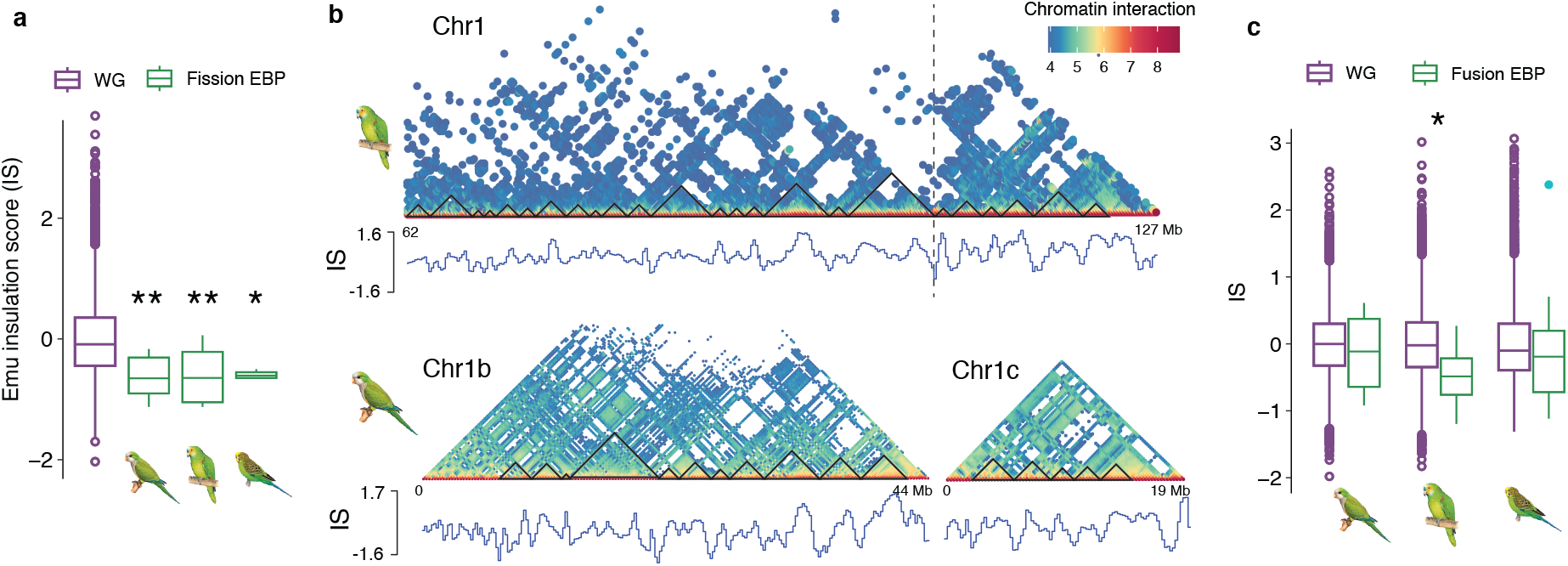
Evolutionary consequence of chromosomal reshuffling. **a)** We compared the insulation scores at the evolutionary breakpoints (EBPs) of fissions between emu and parrot genomes that show significantly lower values compared to the genomic average. * p < 0.05, ** p < 0.01. **b)** one example of chromosomal fission in monk parakeet compared to blue-fronted amazon. The Hi-C matrix was generated at 100-kb resolution. The EBP indicated by a vertical dashed line has the lowest insulation score along the chromosome (only a part of chr1 is shown). **c)** The EBPs of fusion tend to have lower insulation scores compared to the genomic average.

### Chromosomal fusions led to the formation of neo-sex chromosomes

The Hi-C based chromosome-level assemblies and whole-genome alignments (**Fig. 4a**) suggest that chr11 was fused with the ZW sex chromosomes in the ancestor of parrots, and one additional fusion of chr25 with the sex chromosomes specifically occurred in monk parakeet. The FISH experiments using the probes of chicken chr11 and chr25 sequences further validated the chromosomal fusions involving sex chromosomes (**Fig. 4b, Supplementary Fig. S8**). Unlike the scenario of neo-sex chromosome formation in songbirds that involved translocations of parts of chr3-5 (Sigeman, Ponnikas, and Hansson 2020; Gan et al. 2019; Sigeman et al. 2019; Pala et al. 2012; Dierickx et al. 2020), both chr11 and chr25 are microchromosomes, and their entire lengths (23.7 and 2.8 Mb respectively) were added to the sex chromosomes (**Fig. 4a**). To assess whether and to what extent the neo-sex chromosomes are differentiated, we mapped the female sequencing reads to Z chromosomes, with the anticipation that fully differentiated sex chromosomes with divergent Z and W sequences would display reduced coverage. We found the chr11-derived neo-sex chromosome to be hemizygous and fully differentiated but not the chr25-derived one in monk parakeet, consistent with the more recent fusion of chr25. In fact, for the chr25-derived neo-sex chromosome we were not able to assemble the Z- and W-linked sequences separately, likely due to its recent origin with little sequence divergence between them (therefore the coverage is autosome-like, **Fig. 4c**). Re-sequencing data from five additional parrot species (**Supplementary Fig. S9**), and a new chromosome-level assembly of the deep-branching kakapo (Rhie et al. 2020) further support that the fusion of chr11 into sex chromosomes is shared by all parrots.

**Fig. 4.**
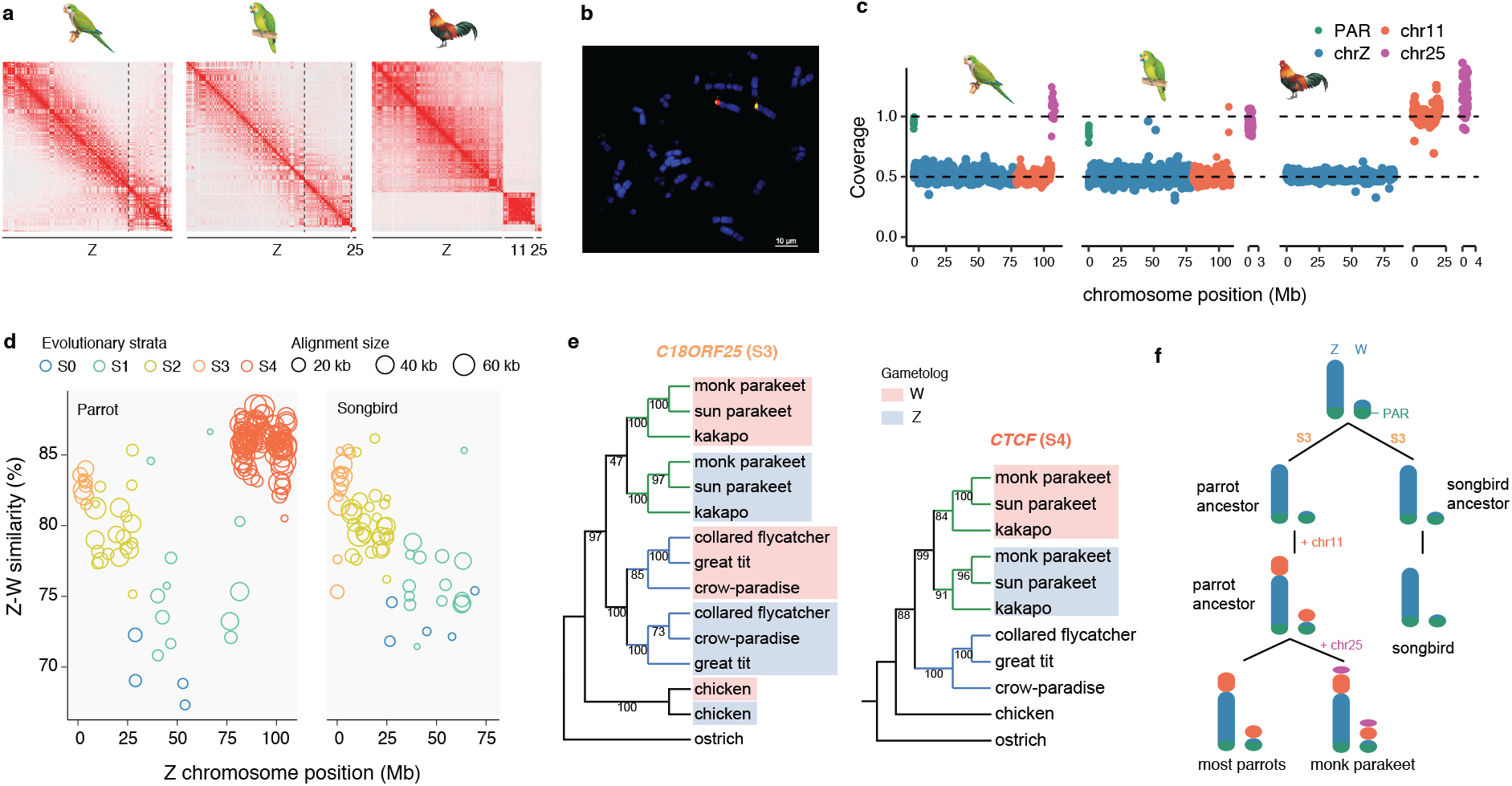
The Evolutionary history of parrot sex chromosomes. **a)** The HiC contact map of chromosomes that are homologous to chicken chr11, chr25 and chrZ. The three chromosomes are presented in three chromosome territories in chicken, but only one in monk parakeet and two in blue-fronted amazon. **b)** The FISH images for the probes of chicken chr25 hybridization in monk parakeet samples. Texas red: CH261-59C21(25q), FITC: CH261-170L23(25p). The FISH result for chicken chr11 probe is shown in the Supplementary Fig. S8. **c)** The female sequencing coverage for the chromosomes homologous to chicken chr11, chr25 and chrZ. In monk parakeet the blue-fronted amazon the coverage of the part of chrZ homologous to chicken chr11 is reduced by half, compared to PARs or autosomes. **d)** Sequence divergence of the Z and W chromosome reveals the pattern of evolutionary strata. Each dot represents a 100-kb sliding window along the Z chromosome. Songbirds (right panel) have four evolutionary strata and parrots (left) have a similar stratum pattern but have one additional stratum (S4) due to the fusion of chr11. **e)** Phylogeny of the Z-W gametologs for a S3 (left panel) and a S4 (right) gametologous gene. Parrot Z- and W-linked gametologs are clustered together respectively, and share a common ancestor, suggesting common origins of S3 and S4 in parrots. **f)** A schematic diagram depicting the evolutionary history of sex chromosome since the common ancestor of songbirds and parrots.

The female coverage patterns also indicate that the old ZW sex chromosomes are fully differentiated, with only a small pseudoautosomal region (PAR) of ~500 kb still recombining between the Z and W chromosomes (**Fig. 4c**). It is known that the sex chromosomes diverge from each other following the arrest of recombination that often occurs in a step-wise manner, forming the so-called evolutionary strata (Cortez et al. 2014; Zhou et al. 2014). By comparing the divergence levels between the Z and W chromosomes, we revealed four evolutionary strata in the old sex chromosome of parrots (**Fig. 4d**), resembling what was found in songbirds (Xu, Auer, et al. 2019), the sister group of parrots (Suh et al. 2011). It was previously demonstrated that all Neoaves (including parrots and songbirds) shared the first three evolutionary strata (S0-S2), with the fourth stratum (S3) often occurring independently among Neoaves lineages (Zhou et al. 2014). Through the phylogenetic analysis of the homologous Z-W gene pairs (gametologs), we confirm that S3 evolved independently in songbirds and parrots (**Fig. 4e**), despite their coincidentally aligned boundaries of PAR and S3 (**Supplementary Fig. S10**).

The chr11-derived neo-sex chromosome formed a single stratum (S4), exhibiting lower divergence than the other evolutionary strata (**Fig. 4d**). This suggests the chr11 was added when S3 was already formed, i.e., the old chromosomes were highly differentiated (**Fig. 4f**). It needs to be noted chr11 (and chr25) were added to the ZW chromosomes at the differentiated end, i.e., not at the recombining pseudoautosomal region (PAR) (**Fig. 4f**), therefore having no impact on the existing PAR. This newly formed S4 retained 16.9% of its original gene content on the neo-W chromosome, compared to only 2.6% on the old W chromosome (**Supplementary Table S4**). The phylogenetic analysis of gametologs from major parrot lineages confirms the origin of the chr11-derived S4 at the ancestor of parrots (**Fig. 4e**).

### Non-adaptive evolution of the neo-sex chromosome

Next we asked whether the newly acquired female-specific neo-W chromosome (S4) evolved a gene repertoire beneficial to females, similar to what had been reported for the neo-Y chromosome of *D. miranda* (Zhou and Bachtrog 2012). The surviving genes (n = 65) on the W-linked S4 all have Z-linked origins, except for a retrogene (*DYNLRB1*) derived from chr12 through LTR-mediated retroposition (**Supplementary Fig. S11a**) (Tan et al. 2016). This intronless retrogene, however, does not seem to be expressed from the W (**Supplementary Fig. S11b**). The expression profile of the S4 Z-W gene-pairs across eight different tissues appears to be highly correlated (**Fig. 5a-b**), suggesting little alteration in gene expression of the W-linked gametologs since their arrival on the W chromosome. However, we detected global down-regulation of the W-linked gametologs relative to the Z-linked gametologs (**Fig. 5a-b**), likely due to the more inactive chromatin environment of the W chromosomes (Xu and Zhou 2020; Liu et al. 2021). Despite that, we identified three genes that are up-regulated across tissues, but only one of them belongs to S4 (**Fig. 5b**). Additionally, W-linked gene loss leads to imbalanced dosage of the proto-sex-linked genes. The avian sex chromosomes are known to have not evolved global dosage compensation (Uebbing et al. 2015; Graves 2014). Consistently, our results suggest that S4 has not evolved a mechanism to fully compensate for the gene dosage across tissues (**Fig. 5c**).

**Fig. 5.**
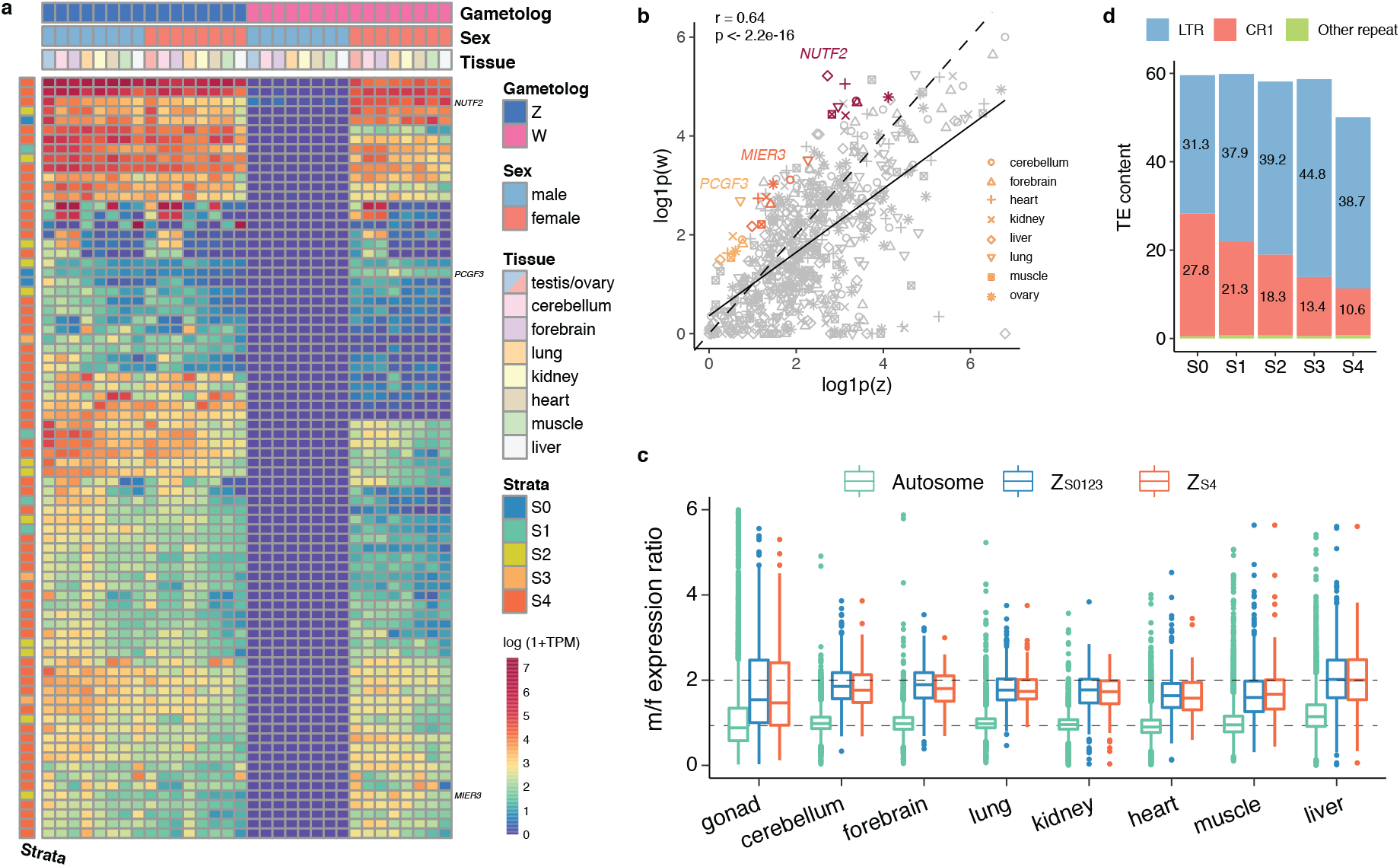
Non-adaptive evolution of the neo-sex chromosome. **a)** A heatmap showing the expression of Z-W gene pairs in male and female tissues in monk parakeet. The W-linked gametologs are present only in females, but their expression profiles are similar to their Z-linked homologs. **b)** Three W-linked gametolog have elevated expression in female tissues relative to the Z-linked homologs. **c)** For both old and new sex chromosomes the expression of the Z-linked genes are lower in female than in male due to dosage imbalance and lack of global dosage compensation. **d)** Temporal dynamics of TE content on the monk parakeet W chromosome divided into five evolutionary strata. LTRs are more abundant in younger strata.

Accompanied with rapid gene loss and down-regulation, the W-linked S4 has already increased the TE content from 12.8% to 50.1% (**Fig. 5d**). The accumulation is mainly driven by LTR element insertions which can be quite long (> 5kb) and tend to remain in their full-length form likely because of the reduced recombination and low efficacy of selection (Leroy et al. 2019; Peona et al. 2020). The young age of S4 also provides a unique window into the temporal dynamics of TE accumulation on the non-recombination sex chromosome, and we demonstrate that LTRs rapidly accumulate on the young strata while CR1 accumulate slower, but over time CR1 gradually increase its proportion as indicated by CR1 densities on older strata (**Fig. 5d**).

### Enlargement of the W chromosome associated with expansions of satellite DNA

The size of the W chromosome in monk parakeet is unusually large and is similar to that of the Z (**Fig. 6a**) (Furo et al. 2017) while in most parrots the W chromosomes are much smaller (Rutkowska, Lagisz, and Nakagawa 2012). Since the chr11 and chr25 were added to both the Z and W chromosomes in monk parakeet, the enlargement of the W chromosome is not due to fusions alone, but likely expansions of repeat sequences (Furo et al. 2017). To identify W-specific repeats, we compared the k-mers from female and male sequencing reads, and found that the top 4 frequent k-mers in females were absent in males (**Fig. 6b**). Those k-mers were derived from a 20-bp satellite (SatW20) which was estimated to have 194,438 copies in the genome, presumably all on the W chromosome. To validate the specificity of SatW20 to the W chromosome, we performed FISH experiments using the monomer of SatW20 as a probe, and confirmed its W-specific binding (**Fig. 6a**).

**Fig. 6.**
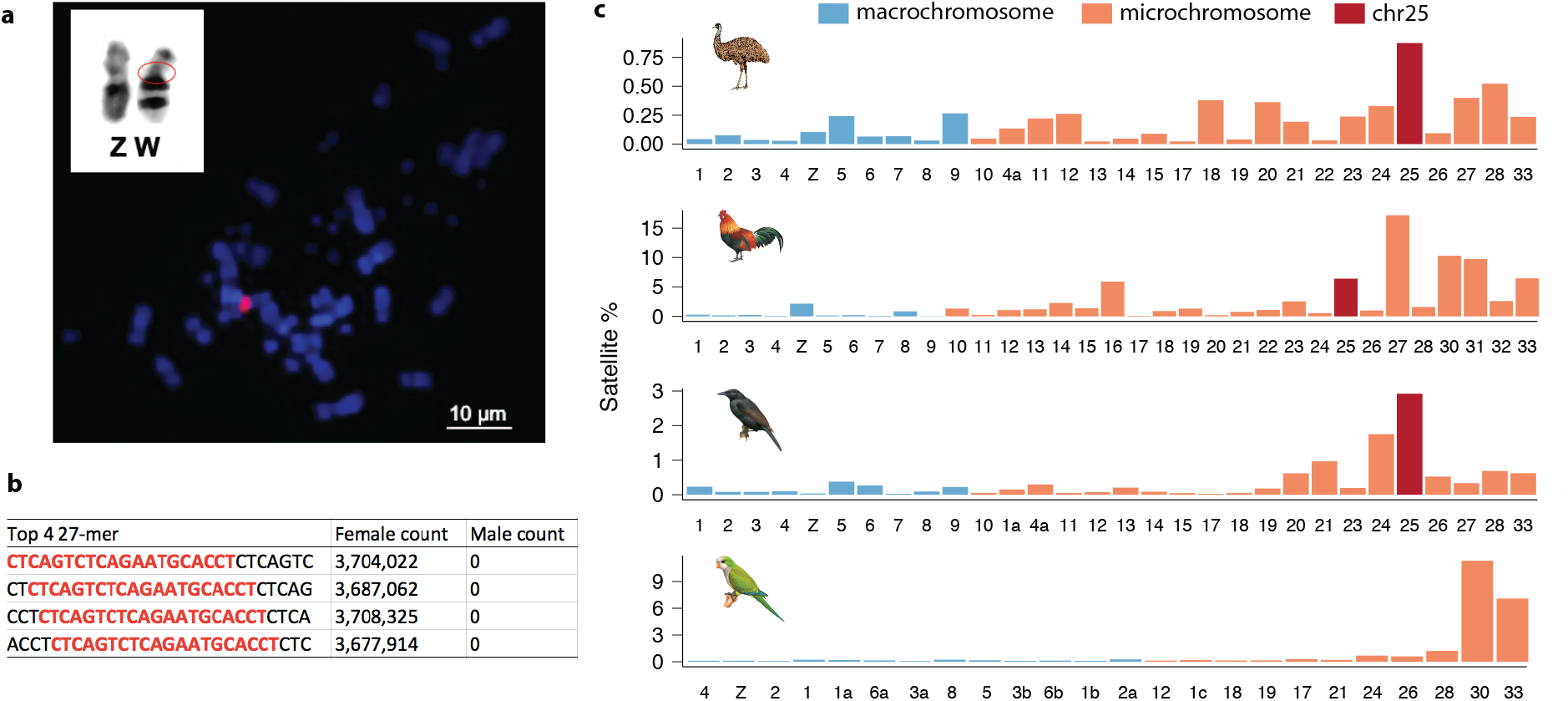
Expansion of satellite DNA contributes to W chromosome enlargement. **a)** G-binding of the ZW sex chromosome and the image of the FISH experiment with the probe of SatW20 (red) for a female cell of monk parakeet **b)** The top four most abundant k-mers (k-mer size 27) in females are absent in male sequencing data. **c)** The proportion of satellite DNA sequences in the chromosomes of four bird genomes assembled with long reads. The chromosome 25 that is fused to the sex chromosomes of monk parakeet is highlighted in dark red.

To unravel the origin of SatW20, we analyzed the composition of repeats of chr11 and chr25 in three other bird genomes assembled with long-reads. In all these birds, chr25 but not chr11 has a considerably large portion of satellite repeats, compared with other chromosomes (**Fig. 6c**). Coincidentally, chr25 was added to monk parakeet specifically. The satellite families of chr25, however, differ among species, indicating rapid turnover of satellite repeats. We were unable to find SatW20 sequences in any of the bird genomes except for the W chromosome of monk parakeet. This suggests that SatW20 had a recent origin following the fusion of chr25 into the ZW chromosome in monk parakeet.

## Conclusions

Through chromosome-level genome assemblies of three parrots, combined with cytogenetic analyses, we uncovered numerous events of chromosomal rearrangements in each of the parrot genomes. The majority of the chromosomal rearrangements are lineage-specific, suggesting karyotypic changes have been regularly taking place in the course of parrot diversification. How chromosomal rearrangements promote speciation of parrots (Kirkpatrick and Barton 2006), however, remains to be investigated. Apart from the frequent chromosomal changes, the parrot genomes have a slightly higher proportion of transposable elements than other birds due to recent CR1 proliferation, and we speculate that recent CR1-psi accumulation may have provided substrates for ectopic recombination events leading to chromosomal rearrangements and the loss of at least some of the 72 conserved genes. Among the lost genes, *ALC1* and *PARP3* are known for their roles in genome stability and DNA double strand repair (Ahel et al. 2009; Day et al. 2017; Mejías-Navarro et al. 2020; Boehler et al. 2011; Beck et al. 2014, 2019; Sellou et al. 2016; Tsuda et al. 2017; Zarkovic et al. 2018; Belousova et al. 2018; Grundy et al. 2016). It has been demonstrated that deletion of *ALC1* impacts chromatin relaxation which is a crucial step in responses to DNA damage (Sellou et al. 2016). Moreover, the depletion of *PARP3* has been shown to delay the repairs of double strand breaks (Boehler et al. 2011; Lindgren et al. 2013; Rodriguez-Vargas, Nguekeu-Zebaze, and Dantzer 2019). Further studies demonstrated that *ALC1* collaborates with *PARP1*, another member of PARP (Poly(ADP-ribose) polymerase) family, on DNA repair (Tsuda et al. 2017; Juhász et al. 2020). This led to our hypothesis that the frequent chromosomal rearrangements were the evolutionary consequences of the loss of both *ALC1* and *PARP3*, which we speculate to have happened through deletions associated with CR1-psi and inversions, respectively. If this were the case, chromosomal rearrangements are perhaps not fixed by natural selection, but rather through genetic drift. Consistent with this notion, we found that the fissions tend to occur at the boundaries of TADs, therefore likely only have slightly deleterious effects on the genomes.

One of the consequences of frequent inter-chromosomal rearrangements is the fusion of sex chromosomes with autosomes, and this has occurred one time at the parrot ancestor and one additional time in monk parakeet. We previously demonstrated that female-specific selection has limited influence on the bird sex-specific chromosomes whose gene content is primarily shaped by purifying selection (Xu and Zhou 2020). Consistently, we did not detect signals of female-specific selection favoring the addition of autosomal female-beneficial genes into the female-specific W chromosomes. In fact, the neo-W chromosomes degenerate and rapidly accumulate TEs, with primarily housekeeping genes surviving, behaving like the old W chromosome (Bellott and Page 2021; Xu and Zhou 2020; Peona et al. 2020); the rapid gene-loss further imposes the challenge of dosage imbalance for female genomes which lack mechanisms of global dosage compensation (Uebbing et al. 2015; Graves 2014). Moreover, the addition of chr25 that is repeat-rich in birds into the sex chromosome in monk parakeet has probably led to the runaway expansion of satellite DNA and heterochromatin on the W chromosome which likely has a deleterious effect for the female genomes (Brown, Nguyen, and Bachtrog 2020; Nguyen and Bachtrog 2020). Together, our study suggests that the frequent chromosomal rearrangements in parrots are are more likely neutral than adaptive, and were probably fixed by genetic drift, involving the role of transposable elements and loss of *ALC1* and *PARP3*.

## Methods

### Long-read sequencing

Parallel genomic DNA extractions were performed on blood from a single female monk parakeet individual from Fuzhou Olsen Agriculture CO.LTD., using the DNeasy Blood & Tissue Kit (QIAGEN, Valencia, CA) following manufacturer’s instructions. The DNA was quantified using the Qubit 2.0 Fluorometer (Thermo Fisher Scientific, Waltham, MA) with its standard protocol. To check for molecular integrity, each DNA was run on the 2200 TapeStation (Agilent Technologies, Santa Clara, CA) following the manufacturer’s protocol. The extracted DNA was used to construct a 20-kb PacBio SMRTbell^TM^ library according to the released protocol from the PacBio Company.

### Re-sequencing

DNA from the feathers of red-and-green macaw (*Ara chloropterus*) and cockatiel (*Nymphicus hollandicus*) were extracted using EasyPure^®^ Genomic DNA Kit (Transgen Biotech, Beijing, China).Then, a paired-end library with an insert size of 350 bp was constructed and sequenced by Annoroad Gene Technology company (Beijing, China) in accordance with the manufacturer’s protocol using the Illumina HiSeq X Ten platform.

### Genome assembly

We used Falcon (pb-assembly v0.0.4) to assemble the long reads into contigs. The haplotigs were removed using the program purge_haplotigs (v1.0.4) (Roach, Schmidt, and Borneman 2018) with the parameter ‘-a 60’. To polish the assembly with long reads, first we mapped the reads to the draft assembly with pbmm2 (0.12.0) which used minimap2 (2.15-r905) for alignment. Then arrow algorism was used for polishing. The mapping-and-polishing process was repeated twice. The short-read sequencing data from the same female individual was used for further polishing. To do so, the reads were mapped to the assembly using BWA-MEM with default parameters. Then pilon (1.22) was used to fix the bases and indels with ‘--minmq 30 --mindepth 20 --diploid’ options. The short-read polishing process was also repeated twice. We used the 3D-DNA pipeline (180114) (Dudchenko et al. 2017) to join the contigs into chromosomes. First, we used the juicer (1.7.6) pipeline (Durand et al. 2016) to map the Hi-C reads against the contigs. Then, we ran the run-asm-pipeline program to do the scaffolding with the options --editor-coarse-resolution 500000 --editor-coarse-region 1000000 --editor-saturation-centile 2 -r 1. The Hi-C contact map based on the draft chromosomal assembly was then visualized in Juicebox which also allowed for manual adjustment of the orientations and order of contigs along the chromosomes. During this process, some misplaced sequences (due to Falcon assembly errors) were cut off from the original contigs and were re-joined to the correct ones.

### Hi-C data analyses

The muscle tissues of monk parakeet and blue-fronted amazon were fixed using formaldehyde for 15 min at a concentration of 1%. The chromatin was cross-linked and digested using the restriction enzyme *MobI*, and then blunt-end-repaired, and tagged with biotin. The DNA was religated with the T4 DNA ligation enzyme. After ligation, formaldehyde crosslinks were reversed and the DNA purified from proteins. Biotin-containing DNA fragments were captured and used for the construction of the Hi-C library. Finally, 350-bp paired-end libraries constructed from DNA were sequenced on the Illumina HiSeq X Ten platform, producing ~80 Gb of sequencing data (**Supplementary Table S2**).

The Hi-C read alignments were performed with the Juicer pipeline. We generated the matrix of Hi-C interaction using the ‘dump’ command of Juicebox at 100-kb resolution with Knight-Ruiz (KR) normalization. We required the interaction count of at least 10 between the pairs of 100 kb interacting windows. This filtered Hi-C matrix was used to visualize the Hi-C contact map. To calculate the insulation scores of TAD boundaries, we used BWA-MEM (-A 1 -B 4 -E 50 -L 0 -t) to map each of the Hi-C read-pairs to the genomes, and generated the Hi-C matrix using the hicBuildMatrix command of HiCExplorer (2.2.1.1). Using the hicFindTADs command, we identified the TAD boundaries and calculated the insulation scores at 200-kb resolution.

### Genome annotation

To annotate the repeat content, we first used RepeatModeler2 (2.0) (Flynn et al. 2020) to predict and classify transposable elements throughout the genome. Tandem repeats were predicted with Tandem Repeats Finder (4.09) (Benson 1999) and the raw results were filtered by the pyTanFinder pipeline (Kirov et al. 2018). The newly predicted families of transposable elements and tandem repeats were then combined with recently curated bird repeat library (including multiple passerines, kakapo, hummingbird, chicken, emu) (Peona et al. 2021) to annotate repeats using RepeatMasker (4.0.7). The repeat-masked genome was then used for the gene model annotation with the MAKER pipeline. The protein sequences from budgerigar, hooded crow and chicken were downloaded from NCBI Refseq database. We mapped the RNA-seq reads from nine different tissues of monk parakeets using HISAT2 (2.1.0) (Kim et al. 2019), and assembled the transcriptomes using StringTie (1.3.3b) with options ‘-m 300 -a 12 -j 5 -c 10’. The gene models were predicted by MAKER based on the alignments of protein sequence and Stingtie transcripts against the genome. To further polish the gene models, we assembled the de novo transcriptome using Trinity (2.8.4) and modified the gene models using the PASA pipeline (v2.4.1) with --stringent_alignment_overlap 30 --gene_overlap 50.

### Transcriptome assembly

For great tit and budgerigar whole genomes were derived male individuals, we downloaded the female RNA-seq data from NCBI SRA to assemble the transcriptomes using Trinity (2.8.4) (Haas et al. 2013), in order to assemble the sequences of W-linked genes. The following options were used in Trinity assembly: “--path_reinforcement_distance 30 --trimmomatic”. We then mapped the RNA-seq reads back to the transcripts with HISAT2 (the above mentioned parameters were used), and the alignments were closely examined in IGV (2.4.3) (Robinson et al. 2011) for potential assembly errors.

### CR1-psi

The draft consensus sequence of CR1-psi, a parrot-specific CR1 element named in this study, was predicted by RepeatModeler2. To curate the consensus sequence, we searched and extracted the homologous sequences in the monk parakeet genome, with an extension of 100 bp to both downstream and upstream directions. The extracted sequences were aligned using MAFFT (7.397) (Katoh and Standley 2013) and the alignment was visualised with Aliview (1.25) (Larsson 2014). We manually inspected the alignments and decided the boundaries of the consensus. The consensus sequence is available in the Supplementary Table S5. We used the parseRM program (Kapusta, Suh, and Feschotte 2017) to estimate the timing of TE insertions at the family level. The phylogeny of known CR1 elements (Suh et al. 2018; Warren et al. 2010) was constructed using FastTree (2.1.11) (Price, Dehal, and Arkin 2010) with default parameters.

### Phylogenomics

We used Last (v1170) (Kiełbasa et al. 2011) to align the genomes of eight birds, including blue-fronted amazon, budgerigar, sun parakeet (Gelabert et al. 2020), kea (Zhang et al. 2014), chicken (Warren et al. 2017), great tit (Laine et al. 2016), paradise crow (Peona et al. 2021), against that of the monk parakeet. The multiple alignment results were then merged with Multiz (v11.2) (Blanchette et al. 2004). A total of 1.63 million four-fold degenerate sites were extracted to construct the phylogenetic tree, using iq-tree with 1000 bootstrap replicates. We then used BASEML (4.9j) to estimate the overall mutation rate with the time calibration on the root node (102 MY). General reversible substitution model and discrete gamma rates were estimated by the maximum likelihood approach under strict clock. The divergence time was then estimated using MCMCtree (4.9j) (Yang and Rannala 2006), with two soft-bound calibration time points: 80-105 MY for the Palaeognathae-Neognathae split and 56-64.5 MY for the Passeriformes-Psittaciformes split (Oliveros et al. 2019).

### Chromosomal rearrangement

We used the MUMmer (4.0.0.beta2) (Marçais et al. 2018) tool nucmer to perform the pairwise whole genome alignment with the parameter “-b 400”. The alignments were filtered to keep the one-to-one best hits using delta-filt from the MUMmer package. Unanchored scaffolds were excluded from the alignments. The alignments were formatted so as to be used by the MCscan pipeline (Tang et al. 2008) for synteny visualization.

### FISH experiment

Chromosome preparations were obtained from fibroblast from feather pulp biopsies of a male and a female of monk parakeet, using standard cell culture and colcemid/hypotonic solution treatment protocols (Furo et al. 2017). Slides were prepared using the air-drying method. FISH with repetitive sequences followed Kubat et al. (2008), and used sequences directly labeled with CY3 as probes. Probes from chicken (GGA) were used to detect segments homologous to chr11 and chr25. We performed FISH experiments using a whole chromosome paint corresponding to chr11, obtained by flow cytometry and labeled with CY3 by DOP-PCR, and BACs (Bacterial Artificial Chromosomes) containing fragments of chr25, labeled with CY3 (short arm) and FITC (long arm), as described previously (O’Connor et al. 2019; Furo et al. 2017). Slides were analyzed with a Zeiss Imager Z2 epifluorescence microscope equipped with a cooled CCD camera and appropriate filters. Images were captured using Axiovision 4.3 (Zeiss), and edited with Photoshop CC.

### Gene expression analysis

We collected nine tissues from three males and three females respectively (**Supplementary Table S2**). The extracted RNA was quantified using the 2100 Agilent Bioanalyzer (Agilent) before library construction. A total quantity of 3 μg RNA per sample was used for paired- end library construction. Sequencing libraries were generated using NEBNext®Ultra™ RNA Library Prep Kit for Illumina® (#E7530L, NEB, USA), following the manufacturer’s recommendations. Briefly, mRNA was purified from total RNA using poly- T oligo- attached magnetic beads (Beckman Coulter). Fragmentation was carried out using divalent cations under elevated temperatures in NEBNext First Strand Synthesis Reaction Buffer (5X). First- strand cDNA was synthesized using random hexamer primer and RNase H. Second- strand cDNA synthesis was subsequently performed using buffer, dNTPs, DNA Polymerase I and RNase H. The library fragments were purified with QIAQuick PCR kits (Qiagen) and eluted with Elution buffer. After terminal repair, poly(A) sequence and adapter were implemented. The resulting cDNA libraries were run on a 2% agarose gel and bands of approximately 250 bp were excised and used for paired- end (2 * 125 bp) sequencing on an Illumina HiSeq X ten platform by Annoroad Gene Technology Co. Ltd.

We use HISAT2 (2.0.4) to map the RNA-seq reads against the genomes with the options -k 4 --max-intronlen 100000 --min-intronlen 30. After sorting the alignments, we used featureCounts (v1.5.2) to count the reads mapped to the annotated gene models. Then TPM (transcripts per million) were calculated to normalize the expression levels for each tissue. The mean TPM values were calculated over the biological replicates.

### Evolutionary strata

The Z and W chromosome sequences were masked for repeats prior to LASTZ (1.04) (Harris 2007) alignment. We used a relaxed parameters (--step=19 --hspthresh=2200 --inner=2000 --ydrop=3400 --gappedthresh=10000) to align the W chromosome to the Z, and alignment chains and nets were further produced to join the syntenic fragmented alignments into longer alignments. Then we filtered out the alignments that are too short (less than 65 bp) or have too low sequence similarity (less than 60%) to reduce the false positive rate. Alignments with unusually high sequence similarity (larger than 96%) were also removed because they might be derived from unmasked simple repeats. We then calculated the sequence similarity between the Z and W over 100-kb sliding windows along the Z chromosome. Because the first (S0) and second (S1) strata are shared by all Neoaves, we used the previously identified sequences of songbird S0 and S1 (Xu et al. 2019) to demarcate the boundaries of S0 and S1 in parrots. We closely examined the Z-W sequence similarity other than the S0/S1 regions, and identified the putative stratum boundary of S2 and S3 at 4.1 Mb. We then used the phylogeny of the Z-W gametologs in the left and right of the putative boundary to test whether they belong to different strata. We used MAFFT (7.427) (Katoh and Standley 2013) to align the coding sequencing of Z-W gametologs and the homologous genes of chicken, ostrich, great tit, collared flycatcher, and paradise crow, with default parameters. We used IQ-TREE (2.0-rc1) (Minh et al. 2020) to perform the phylogenetic analysis, with the substitution model automatically selected. Bootstrapping was repeated 100 times.

## Data availability

The genome assemblies and sequencing data are deposited at NCBI under the accession PRJNA679636. A full list of accession IDs is available in the Supplementary Table S2.

## Code availability

The scripts used in this study have been reposed at Github (https://github.com/lurebgi/monkParakeet).

## Acknowledgement

We thank Zongqin Xu (Tianshen Bird Import and Export Trading Co., Ltd.) for providing the feathers of red-and-green macaw (*Ara chloropterus*) and cockatiel (*Nymphicus hollandicus*), and thank Qi Zhou for comments. L.X. is supported by the Erwin Schrödinger Fellowship (J4477-B) from the Austrian Science Fund (FWF).

